# The line bisection bias stems from left-side underawareness, not from right-side hyperattention

**DOI:** 10.1101/2022.10.21.513001

**Authors:** S. Smaczny, E. Klein, S. Jung, K. Moeller, H.-O. Karnath

## Abstract

It is still a matter of scientific debate whether the line bisection bias frequently observed in patients with spatial neglect is due to attentional underawareness of the left end of the line, attentional hyperattention towards the right end, or a logarithmically compressed perception of the line. To address this question, neglect patients who showed a line bisection bias were administered additional tasks involving horizontal lines (e.g., number line estimation tasks). Their performance was compared to neglect patients not showing a line bisection bias, patients with right hemisphere damage without neglect, and healthy controls. Results indicated that patients with a line bisection bias tended to overestimate lefthand segments when they had to dissect lines into three or four equal parts. This is congruent with both the notions of an underawareness of lefthand segments as well as a logarithmic compression of the line. However, when these patients had to imagine the lines as bounded fraction number lines ranging from 0-1, the results were mixed. When the number lines ranged from 0-10, these patients showed rightward overestimation biases for the numbers 4 and 5. Additionally, all patient groups, but not healthy controls, tended to place number 1 too far to the left and number 9 too far to the right, suggesting a general bias towards endpoints. In sum, this seems more congruent with attentional accounts than a perceptual one. Spatial-numerical associations could be ruled out, as all participants showed a verbal number bisection bias towards smaller numbers (i.e., the ‘left’ of the mental number line). Therefore, these findings seem to indicate that the line bisection bias is most likely due to underawareness of the left end rather than hyperattention towards the right or a logarithmic perception of the line.

## Introduction

Following a (predominantly) right hemispheric stroke, patients often develop spatial neglect, a deficit whereby patients shift their visual orientation towards the ipsilesional side, potentially ignoring objects presented contralesionally. Some patients with spatial neglect show difficulties correctly bisecting a visually presented horizontal line. Instead of indicating the correct midpoint of the line, they typically mark a point that deviates to the right of the actual middle. One interpretation of this deviation has been an asymmetry of lateralised attention allocation (i.e., an attentional bias) between the left and the right hemisphere (Kinsbourne, 1987). According to this account, neglect patients allocate the right end of the line higher attentional weighting than the left end. As line bisection requires equal attentional weighting to both ends of the line to determine the midpoint, those patients misbisect towards the right.

The question remains whether this behavior is due to an unawareness of the contralesional – lefthand (Abe & Ishiai, 2022) or hyperattention towards the ipsilesional – righthand side (Lee et al., 2011). In Abe & Ishiai (2022), patients with neglect used a tablet to bisect lines of different lengths (5, 10, and 20cm), which disappeared directly after bisection. Following this, they had to indicate where either the line’s left or right endpoint had been. They placed the right endpoint accurately and did not differ significantly from healthy control participants. However, they could not reproduce the left endpoint’s correct position reliably, as they tended to mark the same left endpoint position for each line length. In other words, they placed the left endpoint at roughly the same x-coordinate in each trial, irrespective of the left endpoint’s correct position. This observation implies that the right endpoint of a line is correctly represented in patients with neglect. Therefore, the result pattern most likely stems from a misrepresentation of the left end of the line.

This is in contrast with the findings by Lee et al. (2011), who had neurological patients mark either the left quarter (leftmost 25% of the line), the centre, or the right quarter of a line. Neglect patients marked the right quarter significantly more to the right than non-neglect patients and healthy controls, while there was no significant difference in the left quarter mark. As such, these findings suggest hyperattention to the righthand side. However, the authors suggest that these findings may also stem from explicitly asking patients to draw their attention to either the left or right side, which may have provided a top-down ‘override’ of the bottom-up deficit defining neglect (Karnath, 2015). Therefore, it would be more feasible to use a task in which patients are not explicitly cued to one of the endpoints but need to examine the line as a whole.

There are alternative attempts to explain the line bisection bias, according to which line bisection errors are not an attentional phenomenon but of perceptual nature: According to the space anisometry hypothesis, neglect patients perceive the line as logarithmically compressed from one side to the other (Bisiach et al. 2002; Gallace et al., 2008; Pia et al., 2012; Ricci, et al., 2004; Savazzi, et al. 2007). Due to this compression, the right side of the line is perceived to be longer, shifting the subjective midpoint to the right. This hypothesis can be examined when neglect patients mark other proportional elements of the line, not just the centre. If they indeed perceive space anisometrically, the magnitude of their bias should scale logarithmically, depending on which area of the line they are dissecting. This was examined by Gallace et al. (2008): Patients with neglect saw a 16cm line as well as a sample line of 1, 2, 4 or 8 cm. Then they had to divide the 16cm line into accordingly sized segments (i.e., with the 4cm sample they would divide the line into 4cm segments). When patients had to divide the line into two 8cm segments, the left one became significantly longer than the right one. The authors interpreted this finding that the leftward segment was ‘perceived’ to be longer than the rightward segment. There was no effect when patients had to divide the line into smaller segments, that is, dividing the line into quarters or even smaller segments.

Gallace and co-workers (2008) argued that the effect of logarithmic compression only arises when the line is processed ‘globally’ as a whole. For smaller segments, patients focused only on parts of the line at a time, which led them to process the line ‘locally’, thus cancelling out the logarithmic compression of the line when perceived as a whole. Yet, it appears more parsimonious to explain this behaviour via the weighting of fixpoints and using these for orientation: When patients set 8cm marks, both endpoints required an equal attentional weighting to estimate its length (see McIntosh et al., 2005). The endpoint closer to a person’s estimate may then have been weighted more strongly. Accordingly, neglect patients seemed to weight the right endpoint more strongly, biasing their responses to the right. When patients had to divide the line into smaller segments, they only needed to reproduce the distance between one endpoint and the first mark to create subsequent marks, thus overriding the effect of imbalanced weighting of the two endpoints described above.

One way to assess whether neglect patients perceive the line as a whole is to use number line estimation (NLE). In an NLE task, a horizontal line is typically flanked by a number at the start and at the endpoint defining a specific number range. Participants are meant to indicate the spatial location of a certain number on this number line. For example, the left end of the line could be framed as 0, the right end as 10, and participants have to indicate where the number 5 is located on the line. Children (Barth & Paladino, 2011; Dackermann et al., 2018; Jung et al., 2020) and healthy adults (Chesney & Matthews, 2013; Reinert, Huber, Nuerk, & Moeller, 2015) were observed to rely on reference points heavily (i.e., endpoints, midpoints, also referred to as proportion-judgement strategy; Barth & Paladino, 2011, Slusser et al., 2013; but see Booth & Siegler, 2006; Siegler & Opfer, 2003) in such number line estimation tasks.

The use of reference points has also been suggested by McIntosh et al. (2005) for line bisection in neglect patients. This way NLE tasks may serve as an indicator of proportional reasoning instead of purely numerical estimation. That is, participants will orient their response according to certain landmarks on the line, such as the endpoints or the perceived midpoint, irrespective of the number range portrayed by the line. Therefore, the NLE task should be able to differentiate between spatial anisometry, suggesting a deficit in whole line representation, and attentional accounts, suggesting a deficit in the representation of either the left or right end of the line. In particular, assuming that logarithmic compression of the line is the driving factor in line bisection, the difference in rightward bias between neglect patients and controls should be more pronounced in smaller (more leftward) numbers than in larger numbers. However, this should not be the case if line bisection biases are due to the attentional weighting of endpoints. In this case, the rightward bias of smaller numbers should not be more pronounced, or even smaller (see Lee et al., 2011) than for larger numbers.

### Study objectives

The current study addresses whether neglect patients’ line bisection bias is due to hyperattention to the right side, underawareness of the left side, or logarithmic compression of the line, as outlined above. To do so, we administered five different line dissection tasks to (i) a group of patients with neglect displaying a line bisection bias, (ii) one not displaying this bias, (iii) patients with right hemisphere lesions without neglect, and (iv) age-matched healthy control participants.

As a first task, all participants completed a typical *line bisection* task in order to have a baseline line bisection value for the line length of the administered stimuli. In a second task, the *segmentations* task, neglect patients had to subdivide a line into four equal parts (the ‘quarters’ subtask) or into three equal parts (the ‘thirds’ subtask). They had to place all segmentation marks on the same line (other than Lee et al., 2011), ensuring that relative differences between segments could be directly compared for anisometry. Patients could freely choose the points (instead of being provided with a sample line to copy as in Gallace et al. [2008]) to rule out attentional biasing to either end of the line. If line bisection error is indeed due to logarithmic compression of the line, this should be reflected when a line is divided into three or four equally sized parts. The more to the left a mark needs to be placed (i.e., 1/4 compared to 3/4), the larger the patients’ bias to the right should be. A similar pattern should emerge if neglect patients show attentional unawareness of the left end of the line compared to the right side. On the other hand, if hyperattention to the right end of the line leads to bisection deviation, the bias in right marks (i.e., 3/4 or 2/3) should be larger than in left marks (i.e., 1/4 or 1/3). Therefore, the *segmentations* task should be able to differentiate between the hyperattention account and the other two.

In order to further discern whether line bisection bias in neglect patients is due to attentional underawareness or spatial anisometry, participants had to carry out two NLE tasks; one including fractions, and one including whole numbers as stimuli. Yet, in order to keep these tasks comparable to the other tasks and not providing any perceptual cues, flanking numbers at the start and endpoint defining the range of the number line were not written onto the line, as is typical in NLE tasks. Instead, participants were instructed to imagine the line as the number range required for the task (e.g., 0 – 1). If line bisection bias is due to an anisometric perception of the line, this should also influence patients’ estimations of the spatial locations of numbers in NLE (e.g., Barth & Paladino, 2011; Booth & Siegler, 2006; Booth & Siegler, 2008; Chesney & Matthews, 2013; Moeller, et al., 2009; Slusser & Barth, 2017). One would expect the difference in rightward bias between neglect patients and controls to be more pronounced for smaller, and thus more leftward, numbers as compared to larger, and thus more rightward numbers. On the other hand, if line bisection bias is due to effects of attention allocation towards either of the endpoints, this should not necessarily be the case. Framing the line as a number line also has the advantage of make them consider the (number) line as a whole without providing any additional perceptual cues, such as sample lines.

Finally, tasks that involve spatial processing of numbers can potentially introduce spatial-numerical associations (SNAs). That is, when numbers are processed in a spatial dimension, people in left-to-right reading countries tend to represent larger numbers to the right, and smaller numbers to the left, akin to a mentally represented number line (e.g., Dehaene, et al., 1993). Therefore, the NLE tasks (see Fattorini et al., 2016; Pinto et al., 2021a; 2021b) may introduce unwanted rightward biases in neglect patients. In order to rule this out, a verbal number bisection task was carried out (Zorzi et al., 2002). As it has been argued that SNAs need to be activated (Pinto et al., 2021a; 2021b), the number bisection task can examine whether the previous BNLs activate SNAs, whereby patients with neglect should show a systematic rightward bias in the number bisection task only if there is an SNA (Doricchi et al., 2005; Pia et al., 2012; Rossetti et al., 2011, Rotondaro et al., 2015).

## Methods

### Participants

Neurological patients consecutively admitted to the Center of Neurology at Tuebingen University and to the Ermstalklinik in Bad Urach, Germany, were screened for a first-ever right-hemisphere stroke. Patients with visual field defects, a left-sided stroke, patients with diffuse or bilateral brain lesions, patients with tumors, as well as patients in whom MRI or CT scans revealed no obvious lesions were not included. Twenty-one patients with a unilateral, right-sided stroke without visual field defects participated in the study (cf. Table 1). Of these, 11 patients were diagnosed with spatial neglect, and 10 were not (N-). Patients that presented with neglect were further subdivided into patients who showed a line bisection error (NLB+) and those who did not (N+) (see procedure for details). An age-matched (mean, SD) control sample of 32 healthy individuals was also recruited (HC). None of the healthy participants reported any neurological or psychiatric history and all had normal or corrected-to-normal vision. All participants gave written informed consent. The study was carried out in accordance with the Declaration of Helsinki (2013) and was approved by the ethics committee of the University Hospital Tuebingen (Vote 82/2018 BO2).

**Table 1:** Demographic and clinical data of all participants.

|  | HC | N- | N+ | NLB+ |
| --- | --- | --- | --- | --- |
| Age (years) | 66.21 (6.93) | 66.20 (13.98) | 63.2 (14.25) | 73.17 (7.96) |
| Sex F/M (NA) | 16 / 14 (2) | 3 / 7 | 2 / 3 | 1 / 5 |
| Etiology<br>(Ischemia/Hemorrhage) | - | 9 / 1 | 3 / 2 | 3 / 3 |
| Lesion location: | - |  |  |  |
| MCA territory | - | 20% | 100% | 83% |
| MCA & PCA territory | - | 10% | 0% | 0% |
| Basal Ganglia only | - | 40% | 0% | 0% |
| Thalamus only | - | 30% | 0% | 17% |
| Time since lesion (days) | - | 4.5 (4.01) | 62.8 (67.60) | 25 (27.29) |
| Letter Cancellation (CoC) | - | 0.014 (0.021) | 0.534 (0.319) | 0.515 (0.212) |
| Bells Cancellation (CoC) | - | 0.020 (0.039) | 0.579 (0.382) | 0.496 (0.357) |
| Copying task (points) | - | 0.5 (1.27) | 3.6 (2.41) | 3.83 (1.17) |
| Diagnostic line bisection*<br>(% deviation from midpoint) | - | 2.75 (6.10) | 7.41 (6.71) | 34.58 (23.94) |

Data represent mean with SD in parentheses. \*This line bisection task is the line bisection task used in the standard neglect diagnostic battery. It is not the same as the line bisection task considered in our analyses ('task 1').

### Procedure

#### Clinical examination

Stroke patients were administered the following standard diagnostic neglect tests: i) the Letters cancellation (Weintraub & Mesulam, 1985) and Bells cancellation tasks (Gauthier et al, 1989), ii) a copying task (Johannsen & Karnath, 2004), and iii) line bisection (as in Ferber & Karnath, 2001). All tests were presented on a horizontally oriented A4 sheet of paper. For a firm diagnosis of spatial neglect, patients had to fulfill the below criterion in at least two of the four tests.

In the *Letters cancellation* task, 60 target letters ‘A’ are distributed across the sheet of paper amid other distractor letters and patients are asked to cancel out all target letters. The *Bells Cancellation Test* requires patients to identify 35 bell symbols distributed on a field of other symbols. For both the Letter Cancellation Task and the Bells Test, we calculated the Center of Cancellation (CoC) using the procedure and software by Rorden and Karnath (Rorden and Karnath, 2010; www.mricro.com/cancel/). CoC scores greater than 0.08 in the Letter Cancellation Task and the Bells Test indicated neglect behavior (Rorden & Karnath, 2010).

In the *copying task*, patients were asked to copy a complex multi-object scene consisting of four figures (i.e., a fence, a car, a house, and a tree), two each in the left and right half of the test sheet. The following test scores were awarded depending on the correctness of the copied figure: The omission of at least one contralateral feature of each figure was scored 1 point. An additional point was awarded when contralaterally located figures were drawn on the ipsilesional side of the paper sheet. Omitting the complete figure was awarded 2 points. In sum, the maximum score was 8. A score higher than 1 (i.e., > 12.5% omissions) indicated neglect (Johannsen & Karnath, 2004).

The *line bisection task* included 10 lines that were 24cm in length. Lines were presented alternately towards the left or right side of the respective page. Patients were considered to show a bias in line bisection when their estimates deviated from the centre more than 14% towards either side (Ferber & Karnath, 2001).

Additionally, neglect patients were pooled into one group presenting with a line bisection bias (NLB+) and one that did not (N+). Finally, stroke patients were examined with the common neurological confrontation technique for visual field defects.

#### Experimental Tasks

For the experimental tasks, participants were always presented with the same stimuli: 20cm long horizontal lines presented centrally on a horizontally oriented A4 sheet of paper with only one line on a sheet of paper. Participants were explicitly instructed not to measure or, in the case of number lines, not to count in order to derive a solution. Instead, they were told to answer intuitively.

In *task 1*, the *line bisection task*, participants had to bisect lines on 5 separate sheets of paper. In *task 2*, the *segmentations task*, they were required to divide the line into different numbers of equal segments: On five items each they had to divide the lines into either 4 (‘quarters task’) or 3 (‘thirds task’) equally sized segments, in a pseudo-randomised order. To ensure best task performance, participants were explained how many marks they had to set to divide the line into the correct number of segments in case participants did not understand the initial task instructions. In *task 3*, the *fractions task*, participants were asked to imagine the presented line as a number line ranging from 0 to 1. The numbers themselves (i.e., 0 and 1) were not printed on the paper, as flanking numbers are known to affect the dissection of lines (e.g., De Hevia, Girelli, & Vallar, 2006; Fischer, 2001). Participants then had to mark the spatial location of the following fractions 1/4, 1/3, 1/2, 2/4, 2/3, and 3/4 on the number line (each number on an individual sheet of paper), five times each, in a randomized order. This procedure yielded 30 trials in total. *Task 4*, the *whole number task*, was conceptually identical to the previous task, only that the number line was indicated to range from 0-10 instead of from 0-1. Participants then had to indicate the spatial location of the numbers 1, 4, 5, 6, and 9 on the number line. Again, each number had to be estimated 5 times on separate pages, yielding a total of 25 trials.

Finally, participants completed a *verbal number bisection task (task 5*; equivalent to Zorzi, 2002). In the verbal number bisection task, participants were presented with two numbers (ranging from 1 to 22) and were asked to say, as quickly as possible and without calculating, the number bisecting the respective interval – once in ascending order (e.g., ‘what number is exactly between 3 and 7?’ => 5) and once in descending order (e.g., ‘what number is exactly between 7 and 3?’ => 5).

#### Statistical Analyses

Participant responses were measured in cm from the left end of the line. For each participant, mean and SD of responses were calculated for each mark (e.g., mean and SD of marks indicating the number ‘1’ in the whole number task). These means were analysed using the non-parametric Kruskal-Wallis test to compare estimation performance across participant groups. If a task included several measures (e.g., dividing the line into several segments), each segment mark was analysed separately. For example, for the quarters subtask, there was a separate Kruskal-Wallis test for the 1/4, 1/2, and 3/4 marks, respectively. Post-hoc tests were always Benjamini-Hochberg corrected Dunn-Bonferroni tests, and all reported *p*-values represent adjusted *p*-values. For the verbal number bisection task, a single Kruskal-Wallis test was calculated on the mean responses.

## Results

Descriptives for all tasks can be found in table 2.

**Table 2:**
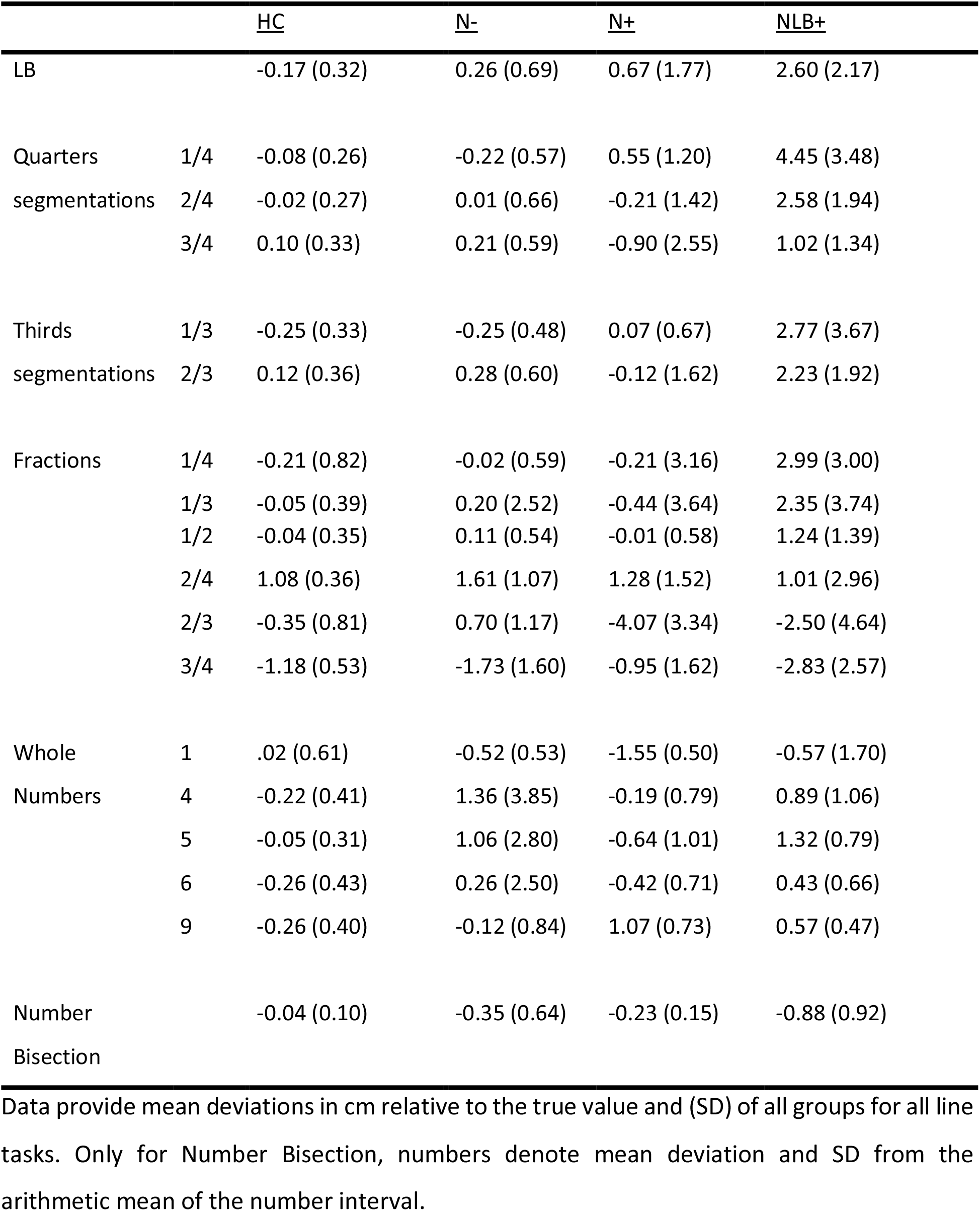
Descriptive values for all groups and all tasks

### Task 1: Line Bisection

One patient from the NLB+ group did not complete this task and is not included in this analysis. Figure 1 illustrates participants’ deviation from the centre of the line in the line bisection task (i.e., *task 1*). As expected, there was at least one significant difference between two of the four groups as indicated by the Kruskall-Wallis test (*H*(3) = 14.728, *p* = .002). Post-hoc tests indicated a significant difference between HC and NLB+ (adj. *p* = .002), whereby patients with a line bisection bias showed a larger deviation from the centre as compared to healthy controls.

**Figure 1:**
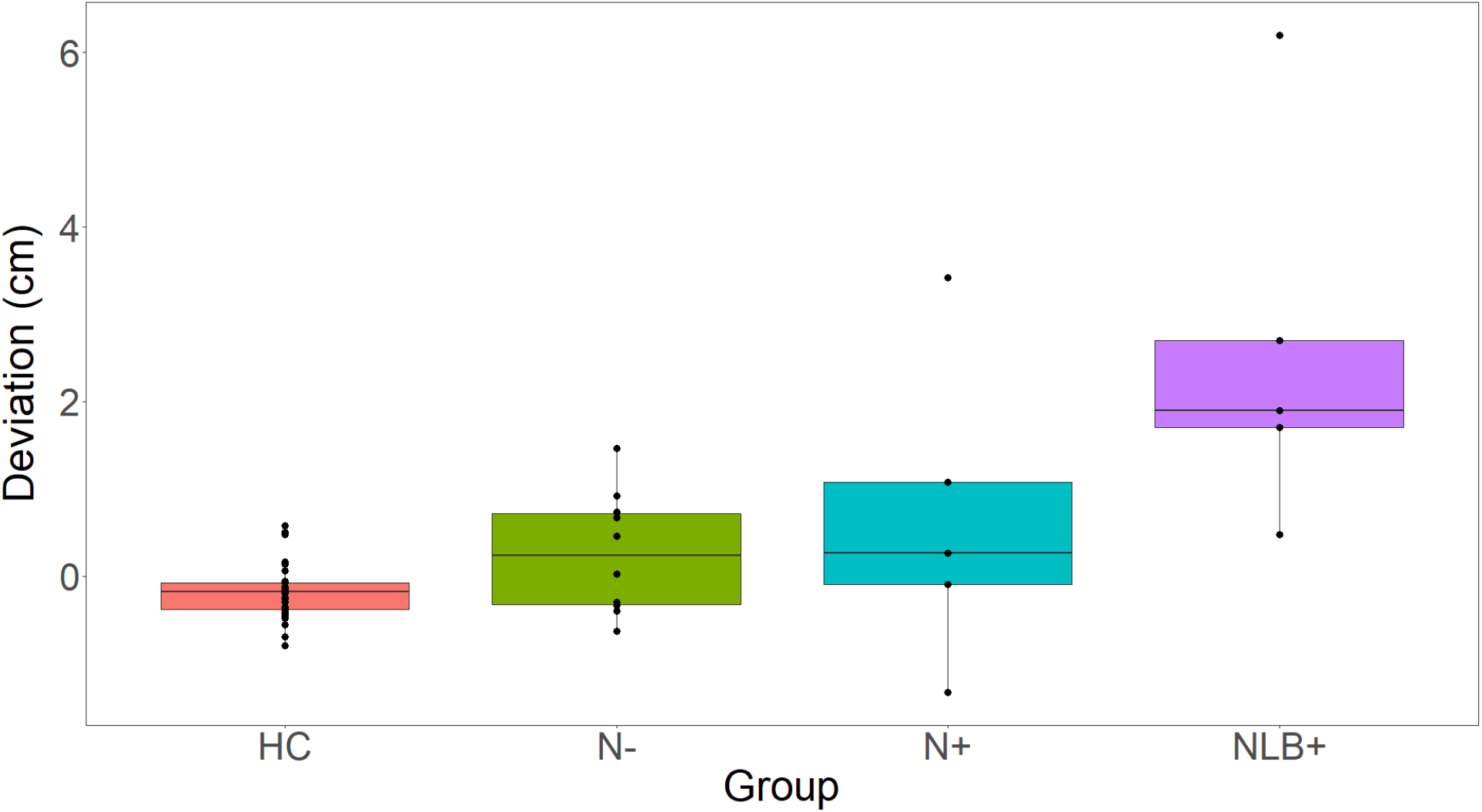
Mean line bisection bias for each group. The y-axis represents participants’ deviation in cm relative to the correct midpoint. Positive values denote a rightward bias. Note that the outlier in group N+ was just under the cut-off of 14% in the diagnostic line bisection task to get classified as NLB+.

### Task 2: Segmentations

In the *segmentations* task, the same lines as used before in *task 1* were presented but now the participants were required to divide the line into either 4 (‘quarters task’) or 3 (‘thirds task’) equally sized parts. Seven patients showed difficulties carrying out this task correctly. Despite explaining thoroughly how many marks they should make to arrive at the correct number of segments, they occasionally set one extra mark. In six of the seven patients, the respective trials were not considered in the analysis; one patient (P15) made too many marks on all trials and was not included in the analysis for this task.

Figure 2 illustrates the difference between participant marks and the correct positions of corresponding marks for the ‘quarters’ subtask; Figure 3 for the ‘thirds’ subtask. Table 3 presents the corresponding results of the accompanying Kruskal-Wallis tests, respectively. There was evidence for significant group differences in both subtasks: For the *quarters* task, the Kruskal-Wallis results indicated significant group differences for the two left lines (i.e., indicating the ¼ and ½ segment of the line). Post-hoc tests revealed a significant difference between the NLB+ and HC group as well as the NLB+ and the N- group for the ¼ mark. Additionally, there was a significant difference between the NLB+ group and all other groups for the ½ line (all adj. *p* < .05). Participants from the NLB+ group placed their marks significantly further to the right than groups HC and N- for the ¼ and all other groups for the ½ mark. In the right line (2/3) of the ‘thirds’ task, groups also differed significantly. NLB+ placed the 2/3 line further to the right than group HC (adj. p < .05).

**Figure 2:**
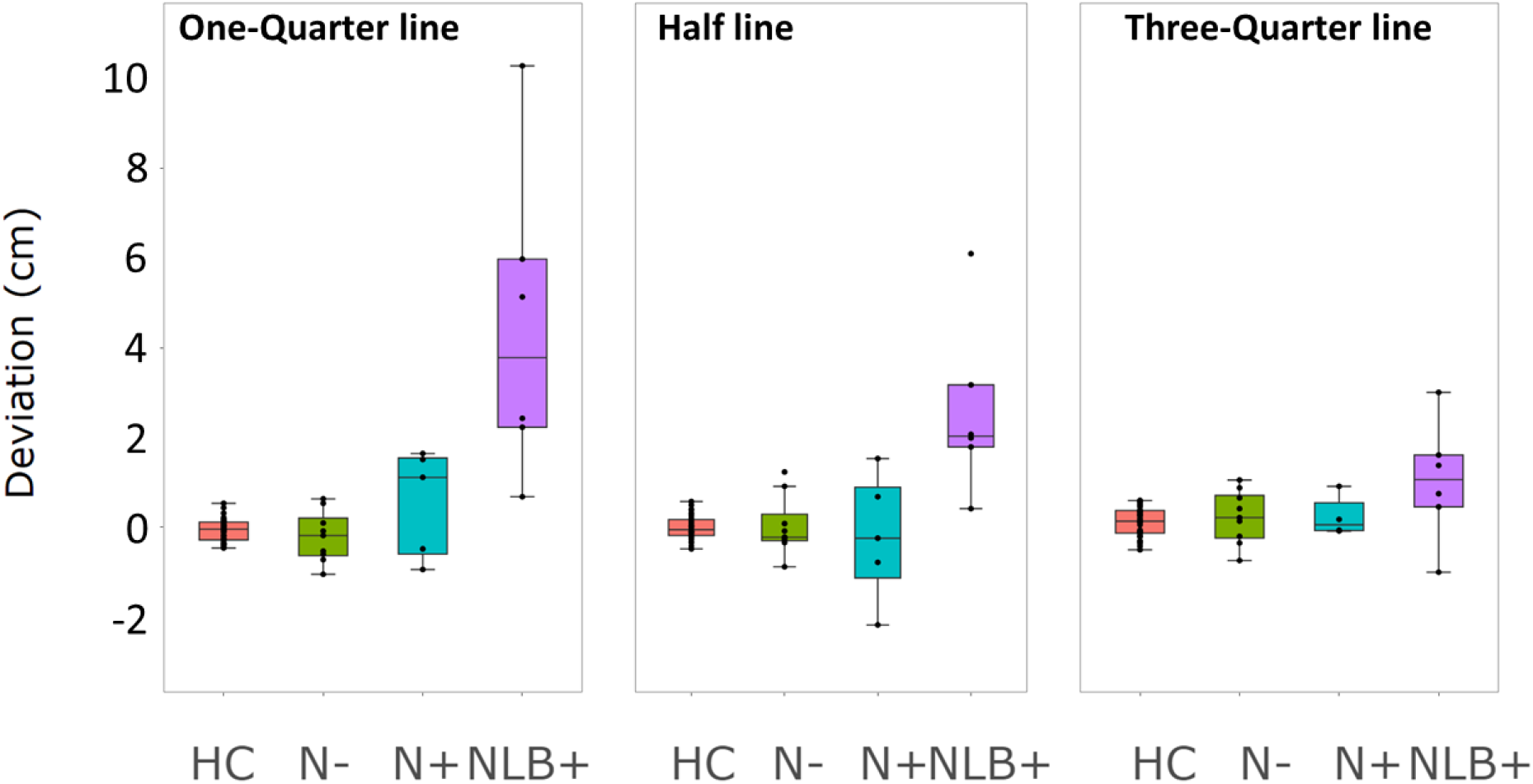
Quarters task. The x-axis reflects groups; the y-axis denotes the mean deviation from the correct response in cm. Black dots present mean responses of individual participants.

**Figure 3:**
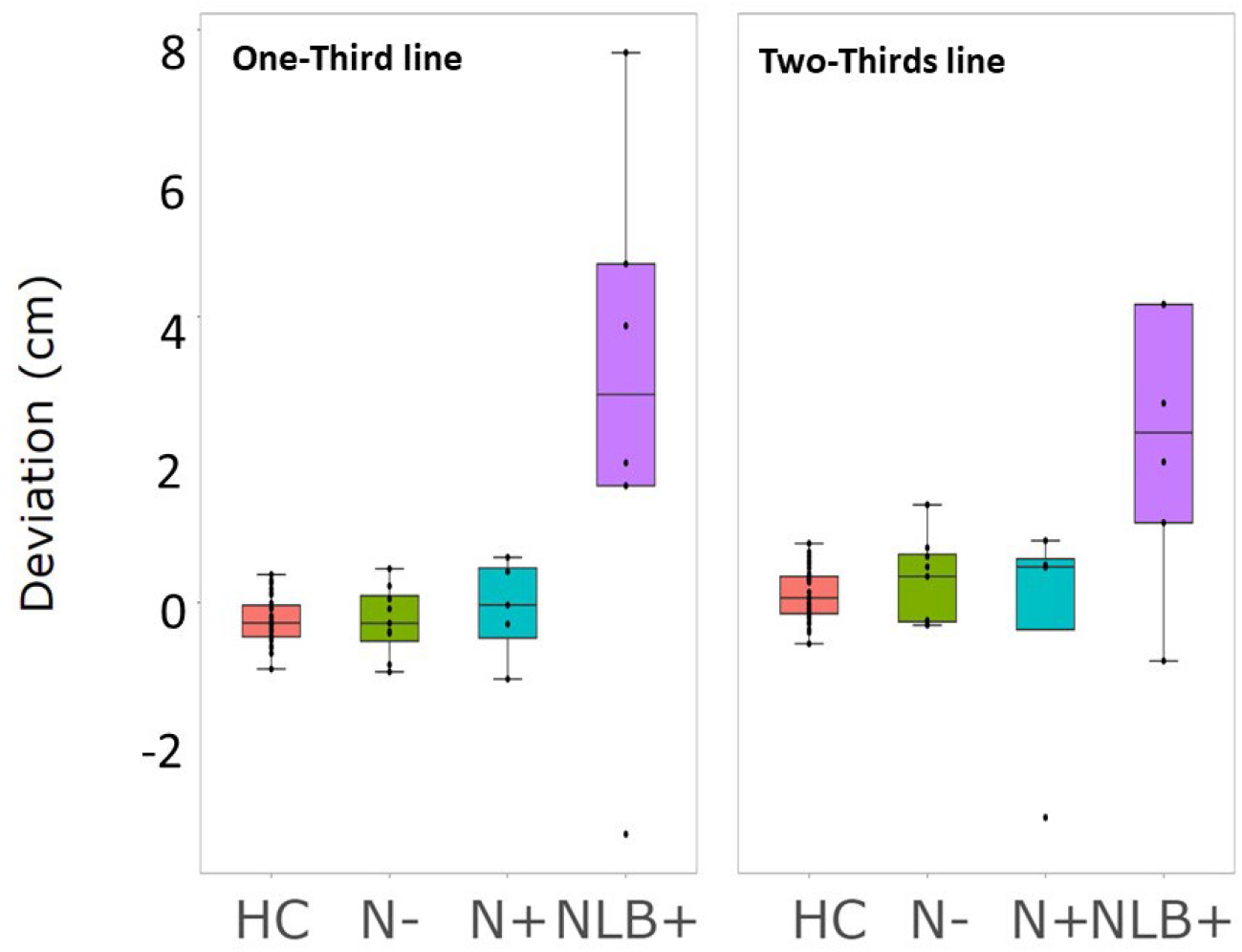
Thirds task. The x-axis reflects groups; the y-axis denotes the mean deviation from the correct response in cm. Black dots present mean responses of individual participants.

**Table 3:** Kruskal-Wallis test results for group differences for each line mark in the ‘quarters’ and ‘thirds’ subtasks.

| Subtask | Position | *H* (df) | *p* |
| --- | --- | --- | --- |
| Quarters | 1/4 | 16.54 (3) | < .001 |
|  | 1/2 | 14.80 (3) | .002 |
|  | 3/4 | 5.86 (3) | .119 |
| Thirds | 1/3 | 7.71 (3) | .052 |
|  | 2/3 | 8.37 (3) | .039 |

Upon closer inspection, in the ‘quarters’ subtask, one N+ (P3) and two NLB+ (P12, P19) patients showed a response pattern indicating that they placed all three lines very close to one another instead of truly dividing the line into equal parts. Also, the same patient from the N+ group (P3) and a different patient from the NLB+ group (P20) behaved very differently from the rest of their groups in the ‘thirds’ subtask (see Figure 3). It seems that these patients could not solve the tasks correctly (see supplementary material for a more in-depth elaboration on these particular patients). Therefore, we assume that these patients may have used a different strategy. Consequentially, we re-analysed data of the *segmentations task*, excluding the respective data (see Table 4). In this re-analysis, Kruskal-Wallis tests for group differences were significant for all items except for 3/4. In particular, we observed a significant difference between the NLB+ group and all other groups for the 1/4 mark. We also observed a significant difference between the NLB+ and the HC as well as the NLB+ and the N- group for the 1/2 mark (all adj. *p* < .05).

**Table 4:** Kruskal-Wallis test results for group differences for each line mark in the ‘quarters’ and ‘thirds’ subtask after exclusions.

| Subtask | Position | $H$ (df) | $P$ |
| --- | --- | --- | --- |
| Quarters | 1/4 | 11.74 (3) | .008 |
|  | 1/2 | 11.33 (3) | .010 |
|  | 3/4 | 2.00 (3) | .572 |
| Thirds | 1/3 | 13.33 (3) | .004 |
|  | 2/3 | 8.95 (3) | .030 |

For the 1/3 mark of the ‘thirds’ subtask, the NLB+ group significantly differed from all other groups (all adj. *p* < .05). For the 2/3 line, there were no significant group differences after correcting for multiple comparisons. Thus, when patients with ‘untypical’ responses were excluded, the NLB+ group showed responses significantly misplaced towards the right compared to other groups for both the ‘quarters’ and the ‘thirds’ subtasks, but only for the leftward marks. Therefore, these results substantiated the results observed with all participants in the *quarters* task, while differing from those including all participants for the *thirds* task. The exclusion of two patients led to significant differences between the NLB+ group with all other groups for the 1/3 line only.

### Task 3: Fractions

In the *fractions* task, participants had to indicate the spatial position of fractions on a number line ranging from 0-1. For this task, two patients had to be excluded from the analyses as they could not complete the task. One was from the NLB+ group (P12), and one from the N- group (P15). Participants’ responses are plotted in Figure 4. Statistical details on group comparisons can be found in Table 5. The only indication for a significant difference was found for 2/3. However, no difference between two groups was significant after correction for multiple testing.

**Figure 4:**
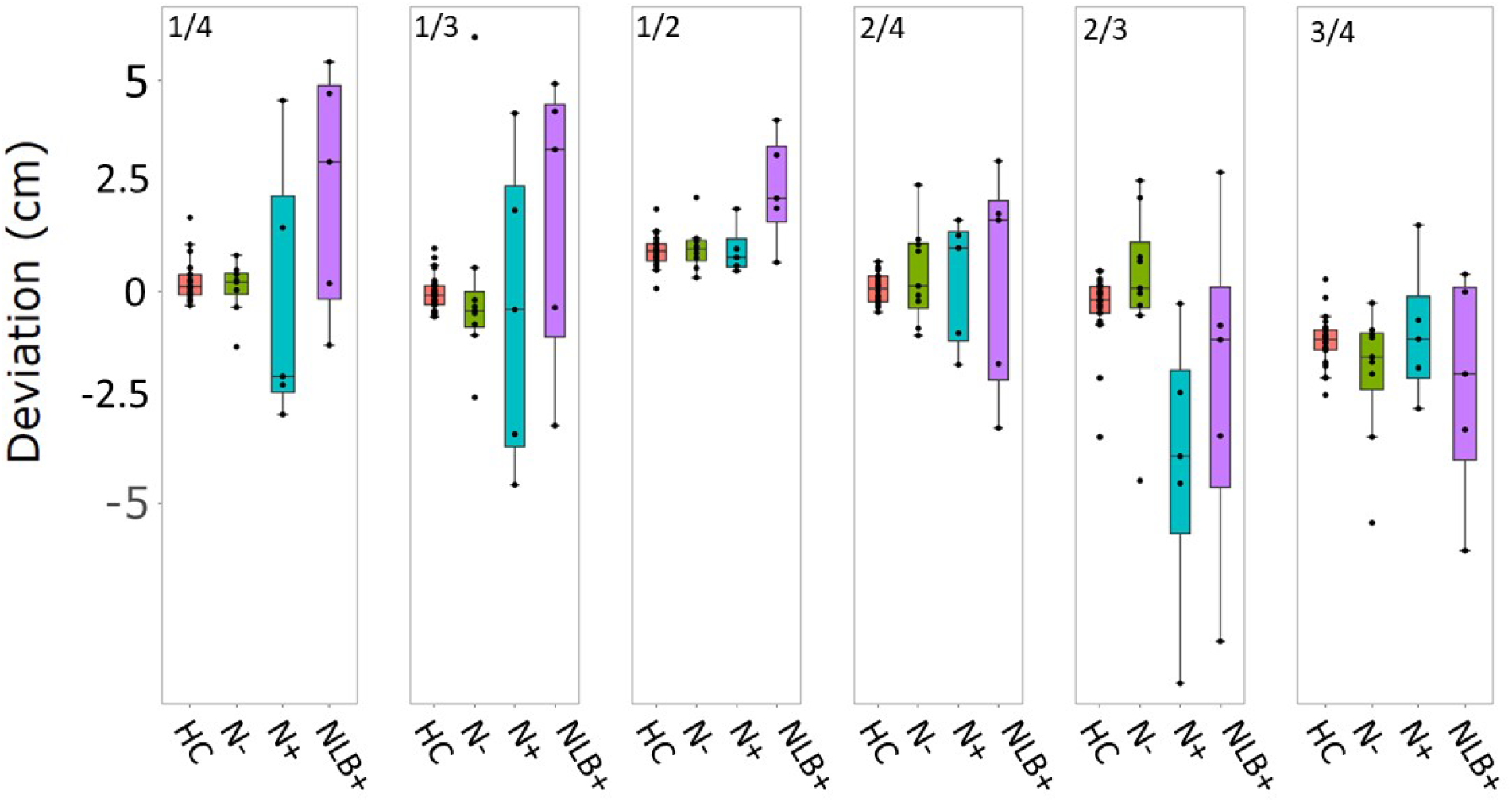
Fractions task. Individual subplots denote group differences in individual marks (i.e., 1/4, 1/3, 1/2, 2/4, 2/3, 3/4). The x-axis indicates group, and the y-axis reflects the relative deviation in cm from the correct position of the respective mark. Black dots indicate individual participants’ mean responses

**Table 5:** Kruskal-Wallis test results for group differences for each line mark in the ‘fractions’ subtask.

| Position | *H* (df) | *p* |
| --- | --- | --- |
| 1/4 | 3.27 (3) | .351 |
| 1/3 | 3.45 (3) | .328 |
| 1/2 | 6.55 (3) | .088 |
| 2/4 | 1.23 (3) | .746 |
| 2/3 | 6.26 (3) | .007 |
| 3/4 | 1.33 (3) | .722 |

### Task 4: Whole numbers

The *whole numbers* task was conceptually identical to *task* 3, only that the number line ranged from 0-10 instead of from 0-1, and participants had to indicate whole numbers. Three patients were not capable of completing the whole number task. Of these, one was the same participant from the NLB+ group who could not carry out the *fractions task* (P12), one was from the N- group (P1), and one was from the N+ group (P3). Therefore, they were not considered in the analyses. Participants’ responses are plotted in Figure 5. Table 6 presents the results of Kruskal-Wallis tests for group comparisons indicating at least one significant group difference for numbers 1, 4, 5, and 9. Post-hoc tests indicated that for number 1, the HC group differed significantly from group N+ (adj. *p* = .003). Additionally, there was a marginally significant difference between the HC group and the NLB+ group after correction for multiple testing (adj. *p* = .063). This indicates that the N+ group placed their marks for the number 1 significantly further to the left compared to the HC group. For number 4, the Kruskal-Wallis test did not indicate any significant differences. For number 5, the NLB+ group differed significantly from all other groups (all adj. *p’s* < .05), placing their mark significantly further to the right than the other groups. There were no significant group differences as identified by post-hoc tests for number 6. For number 9, the NLB+ group differed significantly from the HC group and marginally significantly (adj. *p* = .056) from the N- group. Similarly, the N+ group also differed significantly from the HC and N- groups, whereby N+ and NLB+ placed their marks further to the right than HC and N-.

**Figure 5:**
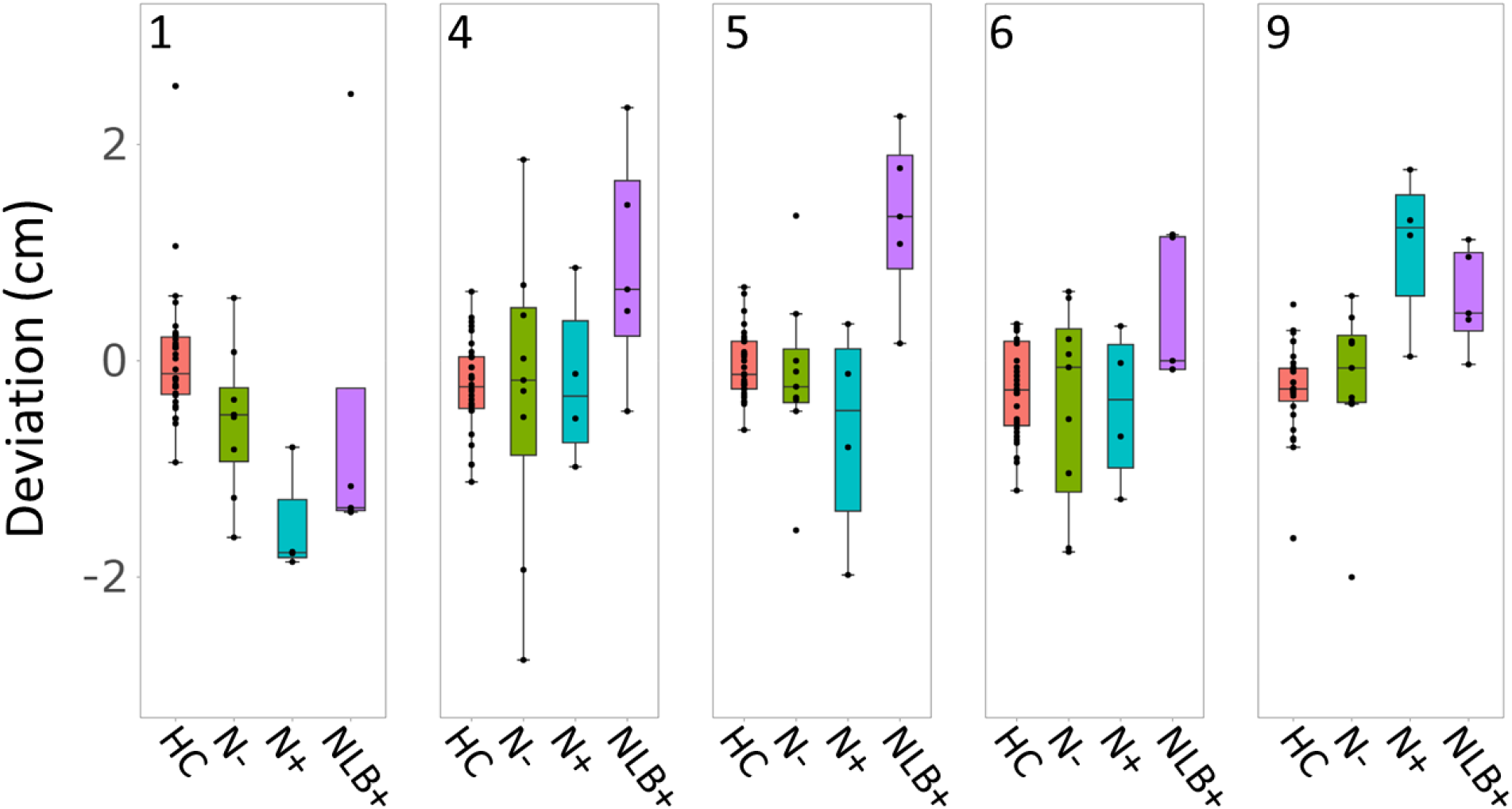
Whole number task. Individual subplots denote group differences in individual marks (i.e., 1, 4, 5, 6, 9). The x-axis indicates group, and the y-axis reflects the relative deviation in cm from the correct position of the respective mark. Black dots indicate individual participants’ mean responses

**Table 6:** Kruskal-Wallis test results for group differences for each line mark in the ‘whole numbers’ task.

| Position | *H* (df) | *p* |
| --- | --- | --- |
| 1 | 17.44 (3) | <.001 |
| 4 | 5.20 (3) | .158 |
| 5 | 11.48 (3) | .009 |
| 6 | 4.10 (3) | .251 |
| 9 | 16.77 (3) | <.001 |

To summarise, for number 1, the N+ group showed a bias to the left compared to healthy controls, for number 5 the NLB+ group showed a bias to the right compared to all other groups, and for number 9 the N+ group showed a bias to the right compared to the HC group and the N- group, while the NLB+ group showed a significant bias to the right compared to the HC group only.

### Task 5: Verbal Number Bisection

In the verbal number bisection task, participants were presented with two numbers (in the range of 1 and 22) and were asked to say, as quickly as possible and without calculating, the number exactly in the middle. Three patients were not able to complete the verbal number bisection task, one of which was again P12 from the NLB+ group, one was P3 from the N+ group, and one was P13 from the N-group. Therefore, these patients were excluded from analyses. Mean patient responses are depicted in Figure 6. A paired samples *t*-test on the entire sample indicated no significant difference between ascending or descending presentation. Therefore, responses were pooled. The HC group’s mean responses were significantly below zero, Wilcoxon Signed Rank test, V = 5, *p* = .025. The same was true for the N- group, V = 0, *p* = .009. There were no significant differences for the groups N+ (V = 0, *p* = .125) and NLB+ (V = 0, *p* = .063). The Kruskal-Wallis test indicated a significant difference between groups (*H*(3) = 25.24, *p* < .001). Closer inspection by post-hoc tests revealed that there was a significant difference between the HC and all other groups (all adj. *p*’s < .05). All patient groups underestimated the midpoint of the given interval significantly more strongly compared to healthy controls. As there were two patients (one from the N- group and one from the NLB+ group) with extreme values, we reran the analysis excluding them. The results were identical (*H*(3) = 22.27, *p* < .001) with the HC group differing significantly from all other patient groups.

**Figure 6:**
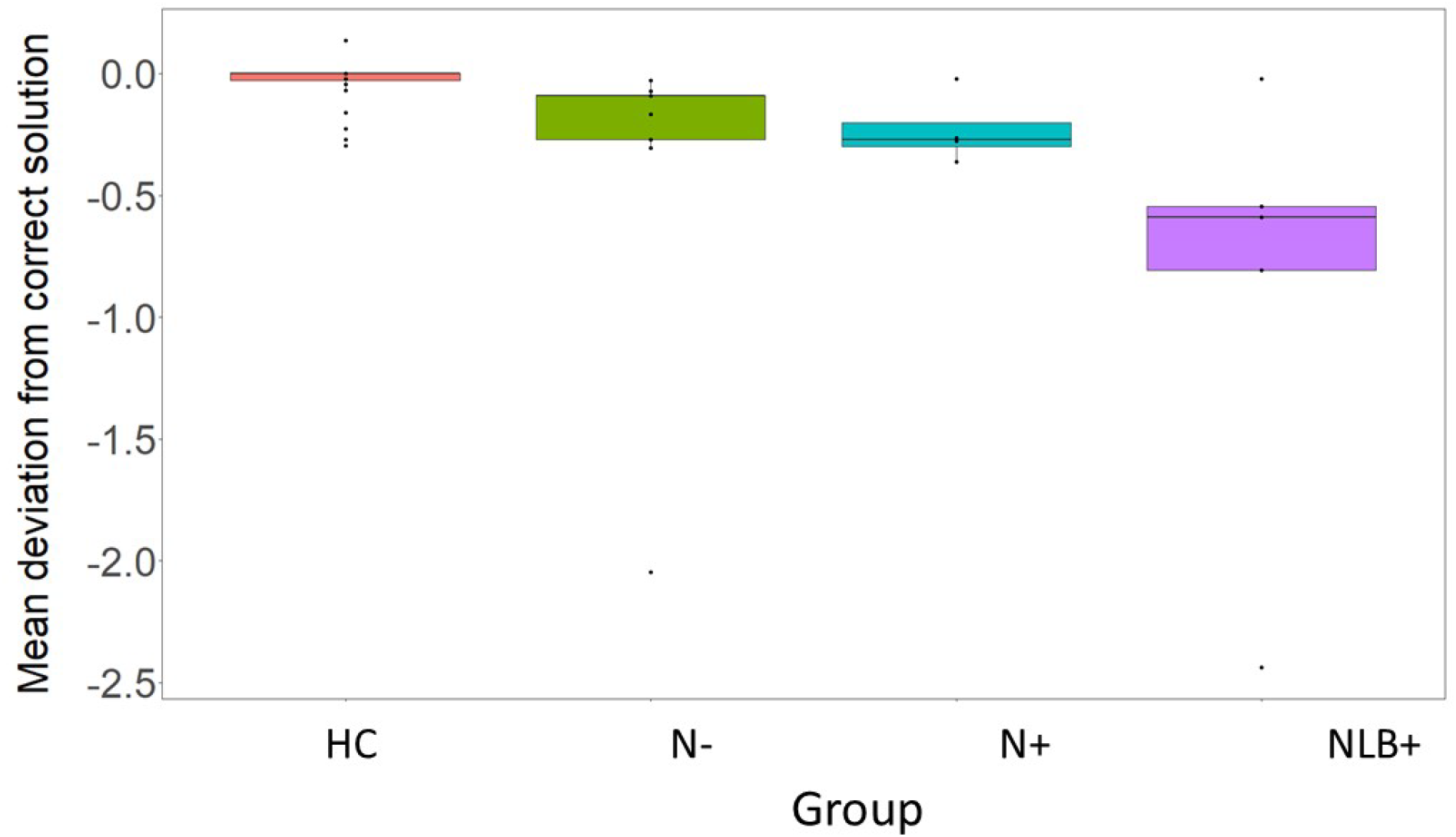
Mean deviation from the correct response for the verbal number bisection deviation for each group in the verbal number bisection task. Dots denote single participants.

## Discussion

In this study, we evaluated whether the typical line bisection bias in spatial neglect can be attributed to either the (i) spatial anisometry hypothesis, which claims that neglect patients perceive the line in a logarithmically compressed way (e.g., Bisiach et al. 2002; Gallace et al., 2008); (ii) hyperattention of the ipsilesional end, whereby patients increase their attention allocation towards the ipsilesional (right) end, thus bisecting ‘towards’ it (e.g., Lee et al., 2011), or (iii) unawareness of the contralesional end, whereby patients seem to be less aware of the left side of the line, and thus bisecting ‘away’ from it (e.g., Abe & Ishiai, 2022).

To this end, patients with neglect presenting with a line bisection bias (NLB+ group), neglect patients not presenting with a line bisection bias (N+ group), patients with right hemisphere damage without neglect (N- group) as well as healthy control participants (HC group) completed different tasks: i) dividing a horizontal line into equally sized segments (quarters or thirds), estimating the spatial location of ii) fractions and iii) whole numbers on the line when framed as number lines (ranging from 0 to 1 and 0 to 10, respectively). Finally, they had to complete a verbal number bisection task in which they had to identify the mean of a given interval. Our expectations were as follows: According to the spatial anisometry hypothesis (i.e., the idea of a logarithmically compressed perception of the line; see Bisiach et al. 2002; Gallace et al., 2008; Pia et al., 2012; Ricci, et al., 2004; Savazzi, et al. 2007), one would expect a larger bisection bias for left marks becoming smaller towards rightward marks in all tasks. In the *segmentations* task this bias should scale from larger to smaller for a quarter, a third, a half, two thirds, three quarters; in the *fractions* task this would be 1/4, 1/3, 1/2, 2/4, 2/3, 3/4; and in the whole numbers task this would be 1, 4, 5, 6, 9. In contrast, when the line bisection bias is based on hyperattention to the right end (Lee et al., 2011), mostly right-sided marks should be set with a bias to the right (i.e., the two thirds / 2/3 and three quarter / 3/4 marks in the *segmentations* task and the *fractions* task and 6 and 9 in the *whole numbers* task). Finally, when underawareness of the left end of the line is the explanation for the line bisection bias, mostly left-sided marks (one quarter and one third in the *segmentations* task, 1/3 and ¼ in the *fractions* task, as well as 1 and 4 in the *whole numbers* task) should be placed with a rightward bias.

Results indicated that in the *segmentations* task, NLB+ patients placed leftward marks (i.e., 1/4, 2/4, 1/3) significantly further to the right compared to the other groups than they did for rightward marks (i.e., 2/3 and 3/4). These results argue against hyperattention towards the right side of the line, contrasting Lee et al. (2011) who reported that neglect patients showed a rightward bias compared to healthy controls in marking the right, but not the left quarter of a line. However, there are some methodological differences between their study and the current one. First, in the current study, patients with a bias in line bisection were examined separately from those without a bias, as previous research has suggested that line bisection is done differently between these groups (e.g., Ferber & Karnath, 2001; Sperber & Karnath, 2016) and has different neural correlates (e.g., Rorden et al., 2006; Verdon et al., 2010; Vossel et al., 2011) than other diagnostic tasks. Second, the experimental tasks were different. In the Lee et al. (2011) study, participants were instructed to mark either the left or the right quarter of the line. Focusing on one-quarter of the line possibly increased the attentional weighting of the closer endpoint. This procedure could have exacerbated a pre-existing bias towards the right endpoint in neglect patients, leading to an inflated bias to the right quarter mark, while possibly balancing out a rightward bias on the left quarter mark.

The emphasis on one or the other endpoint should be less pronounced when participants have to divide the entire line into equal parts, as in the current study. Patients in Lee et al. (2011) also received a training task in which they viewed lines that were pre-divided into four equal parts. This may have influenced their behaviour in the following task, as they could see where they had to place the marks. In the current study, participants were instructed to divide the line into four or three equal segments and were free to set them in whichever order they chose. They were not presented with any pre-divided lines. Therefore, this should have provoked more spontaneous segmentation of the line compared to Lee et al. (2011).

As such, these findings suggest that neglect patients’ bias in line bisection does not seem to be due to hyperattention to the right endpoint. In additional tasks we examined whether this misrepresentation is due to a logarithmic compression of the line (e.g., Gallace et al., 2008) or attentional unawareness of the left side (e.g., Abe and Ishiai, 2022). Importantly, these additional tasks did not comprise visual cues, suggesting that any change in patient behaviour was purely due to a switch in the mental representation of the line. Therefore, behavioural differences between tasks cannot be attributed to changes in the visual perception of the line. The first of these tasks was the *fractions* task, in which participants had to mark the spatial location of fractions when the given line was verbally framed as a number line ranging from 0 to 1. This type of BNL task is typically used to understand the development of magnitude understanding in children (including fractions, e.g., Nuraydin, Stricker, Ugen, Martin, & Schneider, 2023; Siegler & Opfer, 2003), yet a few studies also investigated fraction BNLs in adults (Iuculano & Butterworth, 2011; Siegler & Lortie-Forgues, 2015). In the current study, patient behaviour varied strongly in this task, and there were no significant differences between groups. In general, working with fractions was the most challenging task for participants as indicated by considerable variance within groups. Yet, as these data are very variable, they do not allow a clear-cut interpretation. They do, however, suggest that cognitive reframing of the line may affect both neglect patients’ and control participants’ representation of a line.

When estimating the spatial position of whole numbers, the N+ group placed their marks for 1 significantly closer to the left endpoint than healthy controls, and a marginally significant tendency was observed for NLB+ placing their marks further to the left compared to the HC group. Both the N+ and the NLB+ group placed their marks for 9 closer to the right endpoint than did healthy controls. For numbers 4 and 6 there were no significant group differences. For number 5 the NLB+ group placed their marks further to the right than all other groups. The bias observed for numbers 1 and 9 in the neglect groups (i.e., N+ and NLB+) suggests that when a mark has to be placed close to an endpoint, patients with neglect will have their attention ‘pulled’ towards this endpoint while disregarding the other endpoint.

It has been suggested that the line bisection bias arises due to a limited ability to represent both endpoints simultaneously in neglect patients (e.g., McIntosh et al., 2005; McIntosh & Ishiai, 2022). The current results corroborate this notion. It is also remarkable that neglect patients reversed their rightward bias when having to set their mark close to the left endpoint, as for number 1 in the current task. This may be due to the weighting of endpoints in BNL tasks: Both healthy children (Barth & Paladino, 2011; Dackermann et al., 2018; Jung et al., 2020) and adults (Chesney & Matthews, 2013; Reinert, Huber, Nuerk, & Moeller, 2015) rely heavily on the line endpoints as well as the midpoint as reference points when estimating the spatial location of a given number. For example, when observing participants’ eye movements while carrying out BNL tasks, Reinert et al. (2015) found that participants fixated heavily on these reference points of lines, suggesting proportional judgements to underlie task performance rather than numerical estimation. The current results suggest an exacerbation of this reliance on endpoints as reference points in patients with right hemisphere damage, given their responses to numbers 1 and 9. As such, these findings argue against the spatial anisometry hypothesis (Bisiach et al., 2002; Gallace et al., 2008), as logarithmic compression would suggest that the bias observed for number 1 should be the most extensive rightward bias, at least in the NLB+ group, and decrease as number magnitude of the to-be-estimated number increases.

To counter this, Gallace et al. (2008) suggested that neglect patients may place individual marks differently according to ‘local’ vs. ‘global’ processing. They argued that a visible deviation in marking a line only occurs once individual segments of a line are long enough, as only then the line is processed as a whole. According to this logic, locating numbers 1 and 9 should fall under ‘local’ processing, as they produce segments of only 2cm. In contrast, locating 4, 5, and 6 may fall under ‘global’ processing, as marking these numbers required line segments to be as long (8cm) or longer than the ones used in Gallace et al. (2008). The fact that neglect patients differed from healthy controls and patients without neglect for numbers 1 and 9 contradicts this conclusion.

In contrast, the non-significant group differences for number 6 are, in principal, in line with the notion of a logarithmic compression of the line, as possibly the rightward bias was already ‘too small’ at this point to be found. Yet, the non-significant results for number 4 speak against such a conclusion. Unfortunately, we did not include all numbers between 0 and 10, as this may have enabled more detailed modelling of patient behaviour in the BNL task. Nonetheless, the current results do not support logarithmic compression of the line. Instead, they are compatible with a general deficit in endpoint weighting in neglect patients, which, depending on its severity, may lead to a spatial bias towards the more salient endpoint. In highly severe cases this can manifest itself in a visible line bisection bias (i.e., in group NLB+).

In general, logarithmic compression of lines seems to be unlikely as an explanation for spatial neglect. For example, when neglect patients were required to place LEDs equidistantly along a semicircle, they did not do so in a logarithmically compressed fashion (Karnath & Ferber, 1998). Recent work arguing for the logarithmic compression account has used variants of the Oppel-Kundt Illusion: In these studies, the to-be bisected horizontal line is embedded in a display of several vertical lines that become progressively denser towards one side (Pia et al., 2012; Ricci et al., 2004; Savazzi et al., 2007). The authors of these studies argue that healthy participants’ reaction to this illusion ‘simulates’ the behaviour of neglect patients because healthy participants show a line bisection bias to the right when the vertical lines become denser towards the right side. For neglect patients, the Oppel-Kundt illusion was found to reverse line bisection behaviour when vertical lines became denser towards the left side, leading to a leftward instead of a rightward bias. Similarly, when considering the cognitively framed lines used in the current study, this compression should have worked on the number 1 in the BNL, which it did not.

It has been argued that, under certain circumstances (Pinto et al., 2021a; 2021b), small numbers are associated with the left side of space and large numbers with the right (Dehaene et al., 1993). This assumption has been compared to a continuous ‘mental number line’. Therefore, it could be possible that smaller numbers in the BNL tasks could have induced a leftward bias and larger numbers a rightward one due to spatial-numerical associations. Yet, these SNAs seem to arise only when they are explicitly activated (Pinto et al., 2021a; 2021b). For example, when patients with neglect had to indicate whether a given number was smaller or larger than 5, they showed increased reaction times to smaller numbers when they had to react with their left hand than to larger numbers to which they reacted with their right hand. When they carried out a Go/No-Go task without spatialized responses, there was no effect of SNA (Pinto et al., 2021a). Similarly, when healthy participants were required to react in a Go/No-Go task to numbers or arrows, they reacted quicker when the task instructions explicitly created SNAs (‘press the button when the number is smaller than 5 or when the arrow points to the left’) than when the task instructions did not (‘press the button when the number is smaller than 5 or when you can see an arrow’). This reaction suggests that the association of small numbers with the left and large numbers to the right side of space requires explicit activation of number magnitude information to affect participant responses.

To ensure that our results are not biased through this, we also administered a verbal number bisection task. If SNAs were activated, one should expect neglect patients to overestimate the midpoint of intervals in the verbal number bisection task as compared to healthy controls, as their rightward spatial bias should lead to a corresponding rightward bias on the ‘mental number line’. Yet, there was no systematic ‘rightward’ bias, that means no overestimation of the estimated mean of the respective intervals in neglect patients. This is in line with most literature on verbal number bisection (Doricchi, Guariglia, Gasparini, & Tomaiuolo, 2005; Pia et al., 2012; Rossetti et al., 2011, Rotondaro, Merola, Aiello, Pinto & Doricchi, 2015, but see Zorzi et al., 2002). If anything, NLB+ patients showed a non-significant tendency to produce smaller numbers than other participants, ruling out a lasting association of space and number. Therefore, there is no indication that SNAs affected the BNL tasks and that they were indeed processed as spatial proportions (Barth & Paladino, 2011; Chesney & Matthews, 2013; Dackermann et al., 2018; Jung et al., 2020; Reinert, Huber, Nuerk, & Moeller, 2015).

### Conclusions

The most likely explanation for the results pattern observed in our sample has to do with a weighting of line endpoints: For example, Koyama et al. (1997) examined the bisection behaviour of neglect patients that were grouped into either mild, moderate, or severe neglect on the line bisection task. When they compared their bisection behaviour for lines varying in position and length, the authors found a reliable effect of line length in mild and moderate neglect patients, with longer lines leading to larger bisection biases to the right. In severe neglect patients, the bisections were consistently placed close to the right end of the line, independently of line length. The authors concluded that patients with severe neglect could not shift their attention away from the right endpoint of the line, thus being impaired in ipsilesional attentional disengagement (Posner et al., 1984). This is in line with McIntosh et al. (2005) who argued that patients with severe neglect are not actually capable of bisecting a line; instead, they tend to put attentional weight on the right endpoint and estimate a line midpoint relative to it. Because their attention is so focused on this endpoint, their bisection estimate has a similar distance to the right endpoint at all line lengths. While Lee et al. (2011) conclude that the attentional bias expresses itself through hyperattention to the right side, our results as well as the work of Abe & Ishiai (2022) suggest that it is instead due to a misrepresentation, or underawareness, of the left side of the line. Being ‘unaware’ of the left end of the line also seems more consistent with the more general notion that neglect patients present an underawareness of contralesional space.

## Acknowledgements

This work was supported by the Deutsche Forschungsgemeinschaft (KL 2788/2-1 und KA 1258/24-1). We would like to thank Tamara Matuz, Britta Stammler, and Jasmin Klopfer for their assistance in data collection.

## Notes

https://data.mendeley.com/datasets/tbpb2hj3fn

### Competing Interest Statement

The authors have declared no competing interest.

## References

Barth, H. C., & Paladino, A. M. (2011). The development of numerical estimation: Evidence against a representational shift. Developmental Science, 14(1), 125–135.

Binetti, N., Aiello, M., Merola, S., Bruschini, M., Lecce, F., Macci, E., & Doricchi, F. (2011). Positive correlation in the bisection of long and short horizontal Oppel–Kundt illusory gradients: Implications for the interpretation of the “cross-over” effect in spatial neglect. Cortex, 47(5), 608–616.

Bisiach, E., Neppi-Mòdona, M., & Ricci, R. (2002). 4.1 Space anisometry in unilateral neglect. The cognitive and neural bases of spatial neglect, 145.

Bonato, M., Zorzi, M., & Umiltà, C. (2012). When time is space: Evidence for a mental time line. Neuroscience & Biobehavioral Reviews, 36(10), 2257–2273.

Booth, J. L., & Siegler, R. S. (2006). Developmental and individual differences in pure numerical estimation. Developmental psychology, 42(1), 189.

Booth, J. L., & Siegler, R. S. (2008). Numerical magnitude representations influence arithmetic learning. Child development, 79(4), 1016–1031.

Calabria, M., & Rossetti, Y. (2005). Interference between number processing and line bisection: a methodology. Neuropsychologia, 43(5), 779–783.

Chesney, D. L., & Matthews, P. G. (2013). Knowledge on the line: Manipulating beliefs about the magnitudes of symbolic numbers affects the linearity of line estimation tasks. Psychonomic Bulletin & Review, 20(6), 1146–1153.

Crawford, J. R., & Howell, D. C. (1998). Comparing an individual’s test score against norms derived from small samples. The Clinical Neuropsychologist, 12(4), 482–486.

Dackermann, T., Kroemer, L., Nuerk, H.-C., Moeller, K., & Huber, S. (2018). Influences of presentation format and task instruction on children’s number line estimation. Cognitive Development, 47, 53–62.

De Hevia, M.-D., Girelli, L., Bricolo, E., & Vallar, G. (2008). The representational space of numerical magnitude: Illusions of length. The quarterly journal of experimental psychology, 61(10), 1496–1514.

De Hevia, M.-D., Girelli, L., & Vallar, G. (2006). Numbers and space: a cognitive illusion? Experimental brain research, 168(1-2), 254–264.

De Hevia, M.-D., & Spelke, E. S. (2009). Spontaneous mapping of number and space in adults and young children. Cognition, 110(2), 198–207.

Dehaene, S., Bossini, S., & Giraux, P. (1993). The mental representation of parity and number magnitude. Journal of experimental psychology: General, 122(3), 371.

Doricchi, F., Guariglia, P., Figliozzi, F., Silvetti, M., Bruno, G., & Gasparini, M. (2005). Causes of cross-over in unilateral neglect: between-group comparisons, within-patient dissociations and eye movements. Brain, 128(6), 1386–1406.

Doricchi, F., Guariglia, P., Figliozzi, F., Silvetti, M., Gasparini, M., Merola, S., … Bueti, D. (2008). No reversal of the Oppel–Kundt illusion with short stimuli: confutation of the space anisometry interpretation of neglect and ‘cross-over’in line bisection. Brain, 131(5), e94–e94.

Doricchi, F., Guariglia, P., Gasparini, M., & Tomaiuolo, F. (2005). Dissociation between physical and mental number line bisection in right hemisphere brain damage. Nature neuroscience, 8(12), 1663–1665

Fattorini, E., Pinto, M., Merola, S., D’Onofrio, M., & Doricchi, F. (2016). On the instability and constraints of the interaction between number representation and spatial attention in healthy humans: A concise review of the literature and new experimental evidence. Progress in brain research, 227, 223–256.

Ferber, S., & Karnath, H.-O. (2001). How to assess spatial neglect-line bisection or cancellation tasks? Journal of Clinical and Experimental Neuropsychology, 23(5), 599–607.

Fischer, M. H. (2001). Number processing induces spatial performance biases. Neurology, 57(5), 822–826.

Gallace, A., Imbornone, E., & Vallar, G. (2008). When the whole is more than the sum of the parts: Evidence from visuospatial neglect. Journal of neuropsychology, 2(2), 387–413.

Gebuis, T., & Gevers, W. (2011). Numerosities and space; indeed a cognitive illusion! A reply to de Hevia and Spelke (2009). Cognition, 121(2), 248–252.

Hamdan, N., & Gunderson, E. A. (2017). The number line is a critical spatial-numerical representation: Evidence from a fraction intervention. Developmental psychology, 53(3), 587.

Heilman, K. M., Watson, R. T., & Valenstein, E. (2003). Neglect and related disorders.

Iuculano, T., & Butterworth, B. (2011). Rapid Communication: Understanding the Real Value of Fractions and Decimals. Quarterly Journal of Experimental Psychology, 64(11), 2088–2098.

Jung, S., Roesch, S., Klein, E., Dackermann, T., Heller, J., & Moeller, K. (2020). The strategy matters: Bounded and unbounded number line estimation in secondary school children. Cognitive Development, 53, 100839.

Karnath, H.-O. (1994). Subjective body orientation in neglect and the interactive contribution of neck muscle proprioception and vestibular stimulation. Brain, 117(5), 1001–1012.

Karnath H-O (1994). Spatial limitation of eye movements during ocular exploration of simple line drawings in neglect syndrome. Cortex 30: 319–330.

Karnath, H.-O. (2015). Spatial attention systems in spatial neglect. Neuropsychologia, 75, 61–73.

Karnath, H. O., & Ferber, S. (1998). Is space representation distorted in neglect?. Neuropsychologia, 37(1), 7–15.

Karnath, H.-O., Fetter, M., & Dichgans, J. (1996). Ocular exploration of space as a function of neck proprioceptive and vestibular input—observations in normal subjects and patients with spatial neglect after parietal lesions. Experimental brain research, 109(2), 333–342.

Karnath, H.-O., & Rorden, C. (2012). The anatomy of spatial neglect. Neuropsychologia, 50(6), 1010–1017.

Kinsbourne, M. (1987). Mechanisms of unilateral neglect. In Advances in psychology (Vol. 45, pp. 69–86): Elsevier.

Koyama, Y., Ishiai, S., Seki, K., & Nakayama, T. (1997). Distinct processes in line bisection according to severity of left unilateral spatial neglect. Brain and cognition, 35(2), 271–281.

Lee, B. H., Kwon, S. U., Kwon, J. C., Baek, M. J., Lee, K. H., Kim, G. H., … Na, D. L. (2011). Line quadrisection errors in patients with hemispatial neglect. Neurocase, 17(4), 372–380.

McIntosh, R. D., Ietswaart, M., & Milner, A. D. (2017). Weight and see: Line bisection in neglect reliably measures the allocation of attention, but not the perception of length. Neuropsychologia, 106, 146–158.

McIntosh, R. D., & Ishiai, S. (2022). Endpoints and viewpoints on spatial neglect. Journal of Neuropsychology.

McIntosh, R. D., Schindler, I., Birchall, D., & Milner, A. D. (2005). Weights and measures: A new look at bisection behaviour in neglect. Cognitive Brain Research, 25(3), 833–850.

Moeller, K., Pixner, S., Kaufmann, L., & Nuerk, H.-C. (2009). Children’s early mental number line: Logarithmic or decomposed linear? Journal of Experimental Child Psychology, 103(4), 503–515.

Nuraydin, S., Stricker, J., Ugen, S., Martin, R., & Schneider, M. (2023). The number line estimation task is a valid tool for assessing mathematical achievement: A population-level study with 6484 Luxembourgish ninth-graders. Journal of Experimental Child Psychology, 225, 105521.

Pia, L., Neppi-Mòdona, M., Cremasco, L., Gindri, P., Dal Monte, O., & Folegatti, A. (2012). Functional independence between numerical and visual space: Evidence from right brain-damaged patients. cortex, 48(10), 1351–1358.

Pia, L., Neppi-Mòdona, M., Rosselli, F. B., Muscatello, V., Rosato, R., & Ricci, R. (2012). The Oppel-Kundt illusion is effective in modulating horizontal space representation in humans. Perceptual and motor skills, 115(3), 729–742.

Pinto, M., Pellegrino, M., Lasaponara, S., Scozia, G., D’Onofrio, M., Raffa, G., … & Doricchi, F. (2021a). Number space is made by response space: evidence from left spatial neglect. Neuropsychologia, 154, 107773.

Pinto, M., Pellegrino, M., Marson, F., Lasaponara, S., Cestari, V., D’Onofrio, M., & Doricchi, F. (2021b). How to trigger and keep stable directional Space–Number Associations (SNAs). Cortex, 134, 253–264.

Posner, M. I., Walker, J. A., Friedrich, F. J., & Rafal, R. D. (1984). Effects of parietal injury on covert orienting of attention. Journal of neuroscience, 4(7), 1863–1874.

Ptak, R., Golay, L., Müri, R. M., & Schnider, A. (2009). Looking left with left neglect: The role of spatial attention when active vision selects local image features for fixation. Cortex, 45(10), 1156–1166.

Reinert, R. M., Huber, S., Nuerk, H.-C., & Moeller, K. (2015). Strategies in unbounded number line estimation? Evidence from eye-tracking. Cognitive Processing, 16(1), 359–363.

Ricci, R., Pia, L., & Gindri, P. (2004). Effects of illusory spatial anisometry in unilateral neglect. Experimental brain research, 154(2), 226–237.

Rorden, C., Berger, M. F., & Karnath, H. O. (2006). Disturbed line bisection is associated with posterior brain lesions. Brain research, 1080(1), 17–25.

Rossetti, Y., Jacquin-Courtois, S., Aiello, M., Ishihara, M., Brozzoli, C., & Doricchi, F. (2011). Neglect “around the clock”: Dissociating number and spatial neglect in right brain damage. In Space, time and number in the brain (pp. 149–173). Academic Press.

Rotondaro, F., Merola, S., Aiello, M., Pinto, M., & Doricchi, F. (2015). Dissociation between line bisection and mental-number-line bisection in healthy adults. Neuropsychologia, 75, 565–576.

Salvato, G., Sedda, A., & Bottini, G. (2014). In search of the disappeared half of it: 35 years of studies on representational neglect. Neuropsychology, 28(5), 706.

Savazzi, S., Posteraro, L., Veronesi, G., & Mancini, F. (2007). Rightward and leftward bisection biases in spatial neglect: two sides of the same coin? Brain, 130(8), 2070–2084.

Siegler, R. S., & Lortie-Forgues, H. (2015). Conceptual knowledge of fraction arithmetic. Journal of Educational Psychology, 107(3), 909.

Siegler, R. S., & Pyke, A. A. (2013). Developmental and individual differences in understanding of fractions. Developmental psychology, 49(10), 1994.

Slusser, E., & Barth, H. (2017). Intuitive proportion judgment in number-line estimation: Converging evidence from multiple tasks. Journal of Experimental Child Psychology, 162, 181–198.

Sperber, C., & Karnath, H.-O. (2016). Diagnostic validity of line bisection in the acute phase of stroke. Neuropsychologia, 82, 200–204.

Verdon, V., Schwartz, S., Lovblad, K. O., Hauert, C. A., & Vuilleumier, P. (2010). Neuroanatomy of hemispatial neglect and its functional components: a study using voxel-based lesion-symptom mapping. Brain, 133(3), 880–894.

Vossel, S., Eschenbeck, P., Weiss, P. H., Weidner, R., Saliger, J., Karbe, H., & Fink, G. R. (2011). Visual extinction in relation to visuospatial neglect after right-hemispheric stroke: quantitative assessment and statistical lesion-symptom mapping. Journal of Neurology, Neurosurgery & Psychiatry, 82(8), 862–868.

Zorzi, M., Priftis, K., & Umiltà, C. (2002). Neglect disrupts the mental number line. Nature, 417(6885), 138–139.

