## Supplemental figures for "The line bisection bias stems from left-side underawareness, not from right-side hyperattention"

### Supplementary

Figure S1 presents mean responses for one individual patient in the “quarters” subtask who attempted the task but showed highly untypical responses compared to the rest of their group. Group N+ generally was able to divide the line into 4 equal parts, only patient #3 placed all three lines very close to one another, and all of them into the left half of the line (see also Figure 2 of the main manuscript). This suggests that patient #3 was not able to correctly carry out this task. For this reason, we excluded patient #3 for a second, additional analysis.

Figure S1: A comparison of patient #3 with the rest of group N+ in the “quarters” subtask

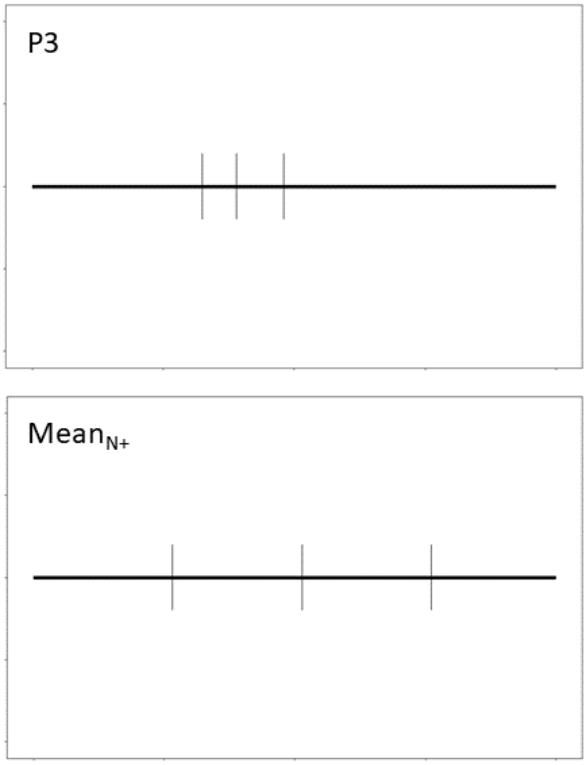


Such highly untypical response patterns could also be found for two members of group NLB+ in the “quarters” subtask (cf. Figure S2). While showing great variation, Group NLB+ generally showed behavior indicating an attempt to divide the line into 3 equal parts, only that their responses were all skewed towards the right, as if they were unaware of the left end of the line. On the other hand, patients #12 and #19 just placed all lines very close to one another, suggesting that they were not able to carry out the task correctly. Therefore, we excluded them for a second, additional analysis of the “quarters” task.

Figure S2: A comparison of patients #12 and #19 with the rest of the NLB group

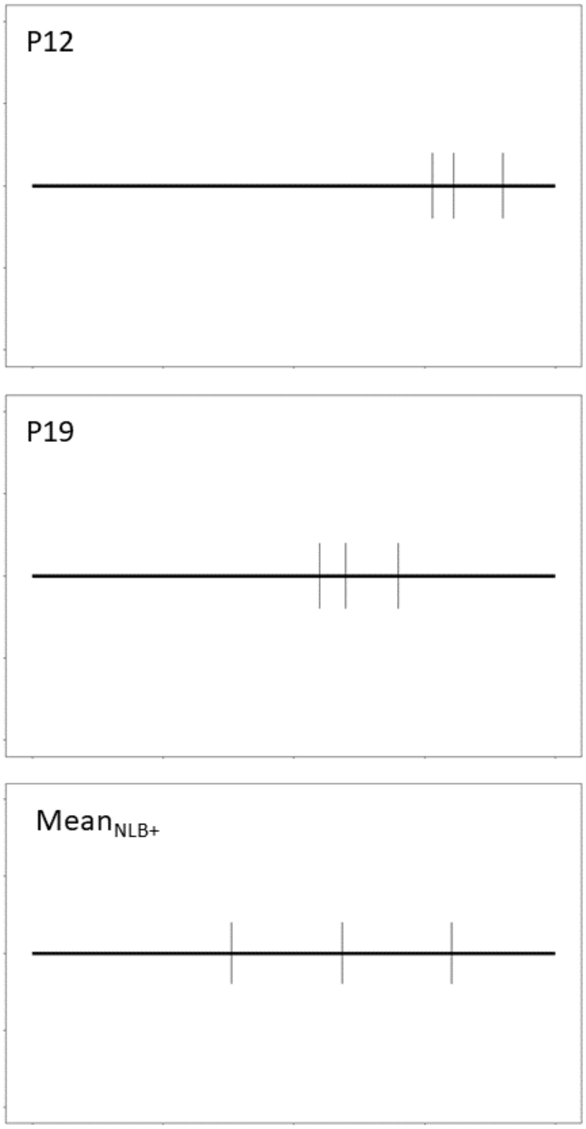


Similar issues arose for the “thirds” subtask. Again, patient #3 placed both marks very close to one another, while the rest of group N+ showed no pronounced difficulties dividing the line (Figure S3). Interestingly, patient #20 of group NLB+ showed highly unusual behaviour in the “thirds” subtask, placing their left mark the most extremely to the left and their right mark the most extremely to the right (Figure S4). Therefore, patient #20 was also excluded from the analysis of the “thirds” task.

Figure S3: A comparison of patient 3 with the rest of group N+ in the “thirds” subtask

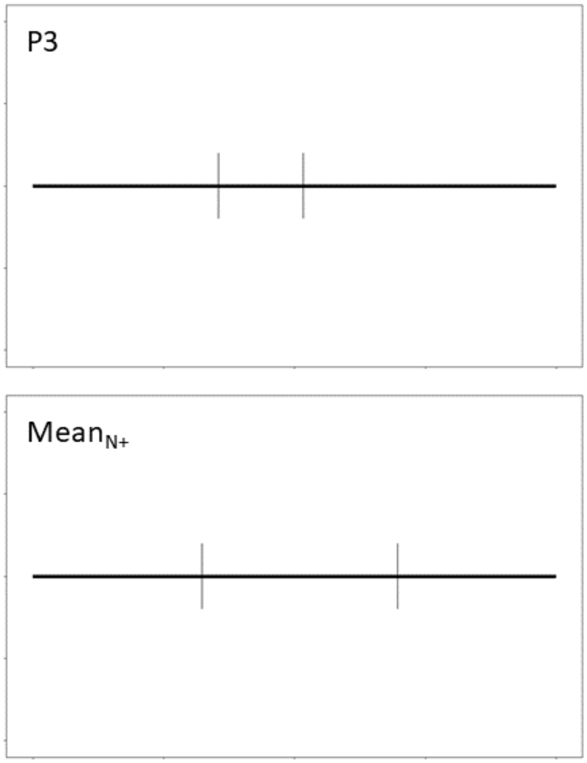

Figure S4: A comparison of patient #20 with the rest of group NLB+ in the “thirds” subtask

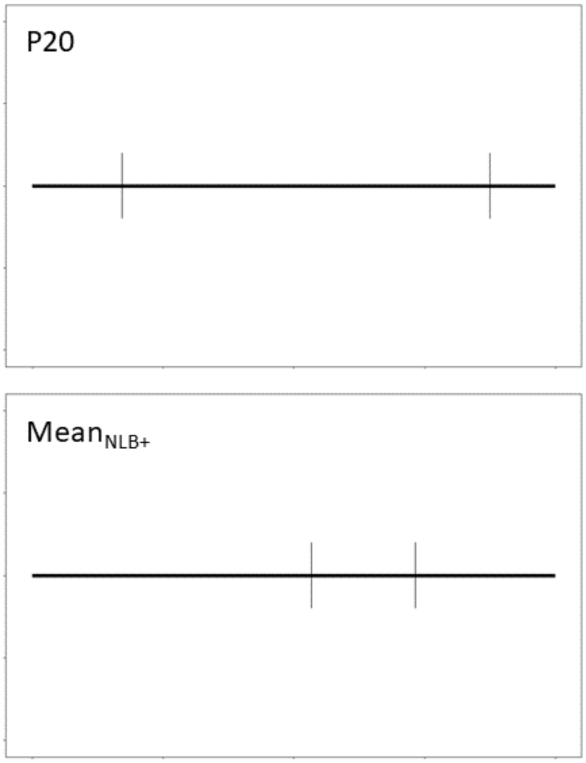
